# Comprehensive Identification of Fim-Mediated Inversions in Uropathogenic *Escherichia coli* with Structural Variation Detection Using Relative Entropy

**DOI:** 10.1101/494401

**Authors:** Colin W. Russell, Rashmi Sukumaran, Lu Ting Liow, Balamurugan Periaswamy, Shazmina Rafee, Yuemin C. Chee, Swaine L. Chen

## Abstract

Most urinary tract infections (UTIs) are caused by uropathogenic *Escherichia coli* (UPEC), which depend on an extracellular organelle (Type 1 pili) for adherence to bladder cells during infection. Type 1 pilus expression is partially regulated by inversion of a piece of DNA referred to as *fimS*, which contains the promoter for the *fim* operon encoding Type 1 pili. *fimS* inversion is regulated by up to five recombinases collectively known as Fim recombinases. These Fim recombinases are currently known to regulate two other switches: the *ipuS* and *hyxS* switches. A long-standing question has been whether the Fim recombinases regulate the inversion of other switches, perhaps to coordinate expression for adhesion or virulence. We answered this question using whole genome sequencing with a newly developed algorithm (Structural Variation detection using Relative Entropy, SVRE) for calling structural variations using paired-end short read sequencing. SVRE identified all of the previously known switches, refining the specificity of which recombinases act at which switches. Strikingly, we found no new inversions that were mediated by the Fim recombinases. We conclude that the Fim recombinases are each highly specific for a small number of switches. We hypothesize that the unlinked Fim recombinases have been recruited to regulate *fimS*, and *fimS* only, as a secondary locus; this further implies that regulation of Type 1 pilus expression (and its role in gastrointestinal and/or genitourinary colonization) is important enough, on its own, to influence the evolution and maintenance of multiple additional genes within the accessory genome of *E. coli*.

**Importance:** UTIs are a common ailment that affects more than half of all women during their lifetime. The leading cause of UTIs is UPEC, which rely on Type 1 pili to colonize and persist within the
bladder during infection. The regulation of Type 1 pili is remarkable for an epigenetic mechanism in which a section of DNA containing a promoter is inverted. The inversion mechanism relies on what are thought to be dedicated recombinase genes; however, the full repertoire for these recombinases is not known. We show here that there are no additional targets beyond those already identified for the recombinases in the entire genome of two UPEC strains, arguing that Type 1 pilus expression itself is the driving evolutionary force for the presence of these recombinase genes. This further suggests that targeting the Type 1 pilus is a rational alternative non-antibiotic strategy for the treatment of UTI.

## Introduction

Uropathogenic *Escherichia coli* (UPEC) are the primary cause of urinary tract infections (UTIs) (1, 2), which are estimated to affect more than half of all women during their lifetime (3). The total annual cost of community-acquired and nosocomial UTIs in the United States was estimated to be $2 billion in 1995 (3). Although UTIs have traditionally been effectively treated with antibiotics, in some patients UTIs recur despite apparently appropriate antibiotic therapy and sterilization of the urine (4). Furthermore, UTIs are the first or second most common indication for antibiotic therapy (5, 6), making them a major contributor to rising antibiotic resistance rates (7). Therefore, substantial effort has been devoted to studying the molecular mechanisms by which UPEC cause UTI in the service of developing alternative preventive and therapeutic strategies (2, 8-11).

One of the major successes in UTI research has been the recognition of the importance of Type 1 pili for causing UTI (12-14). Type 1 pili, encoded by the *fim* operon, are hair-like, multiprotein structures that extend from the outer membrane and terminate in the adhesin protein FimH (15-17). FimH binds to mannose residues on glycosylated bladder surface proteins such as uroplakin protein UPIa (18) and α3β1 integrin heterodimers (19). Adhesion to the bladder epithelium can lead to internalization of the bacteria into host cells and formation of intracellular bacterial communities (IBCs) (20-23). Bacteria in IBCs are protected from the immune response and antibiotic treatment, and can later escape from the host cells to cause recurrent infection (24, 25). Therefore, Type 1 pili directly contribute both to the initiation of infection and to intracellular persistence. Several new strategies have focused on blocking the function of Type 1 pili by small molecule inhibition or vaccination (26, 27).

The pilus structural proteins (including the FimH adhesin) and the chaperone-usher proteins that mediate pilus biogenesis are encoded within the *fimAICDFGH* operon (15, 16). Regulation of Type 1 pili expression centers on the epigenetic alteration of the *fim* operon promoter, which is located within the invertible *fim* switch *fimS* (28, 29). When *fimS* is in the ON orientation, the promoter is positioned to transcribe the *fim* genes and Type 1 pili may be synthesized. In contrast, when the *fimS* promoter is in the OFF orientation, bacteria do not produce Type 1 pili.

Switching of *fimS* from one state to another is regulated by recombinases which bind to inverted repeat (IR) sequences that flank the switch. Two recombinases, FimB and FimE, are encoded by genes that are genetically linked to the *fim* operon and *fimS* switch (30). Other known recombinases acting at *fimS* include the genetically unlinked IpuA and FimX (30-32). Interestingly, both the linked and unlinked Fim recombinases are also able to mediate the inversion of other switches. The *hyxS* switch is inverted by FimX (33), while *ipuS* was shown to be inverted by FimE, FimX, IpuA, and IpuB (but not FimB) (34). Like *fimS*, inversion of *hyxS* and *ipuS* appears to regulate downstream gene expression, but the full importance of these genes in pathogenesis is still not clear.

An open question in the field has been whether the Fim recombinases are utilized in the regulation of other, still unknown, switches, and whether such switches may be related to pathogenesis. To search for novel invertible elements, we developed an algorithm named Structural Variation detection using Relative Entropy (SVRE) to detect genomic structural variations (SVs) in whole genome sequencing data. We applied SVRE to uropathogenic strains overexpressing each Fim recombinase. In addition to the known inversions at *fimS*, *hyxS*, and *ipuS*, SVRE detected several SVs that were recombinase-independent. Importantly, no new invertible switches were found, indicating that *fimS* is inverted by several recombinases that regulate little else, suggesting that tuning of Type 1 pilus expression is of strong evolutionary importance.

## Results

### Development of SVRE

Invertible sequences like *fimS* are one class of SV, which also includes deletions, duplications, translocations, and more complex rearrangements. Several programs have been developed to call SVs from whole genome sequencing data. One primary strategy for SV detection is to identify paired-end reads with unusual mapping patterns. Generation of DNA libraries for next-generation sequencing typically includes a size selection step that restricts the physical size of the DNA fragments that are carried forward for sequencing. When mapped to an ideal reference genome, the distance between paired-end reads should reflect this length. Additionally, the reads should map to opposite strands of the genome. Paired-end reads with an appropriate mapping distance and read orientation are termed “concordant” reads. In contrast, in the presence of an SV in the input DNA relative to the reference genome, paired-end reads associated with the SV map at a distance or orientation that differs from this expectation; these reads are called “discordant” reads.

We developed SVRE, an algorithm that detects SVs by analyzing the distribution of mapping distances in segments of the genome. When reads span an SV, the local mapping distances for these reads should follow a different distribution based on the type of SV; the difference in distribution is generated by discordant reads. In the case of an invertible element like *fimS*, the genomic material used for sequencing may contain a mixture of both orientations (Figure 1A). Reads derived from the invertible element will map to the reference genome differently depending on the orientation of the element. If the orientation is the same as the reference, the reads will align with the expected mapping distance to opposite strands (the gray arrows in Figure 1A). However, if the orientation is reversed, the paired-end reads will map to the same strand and with a mapping distance different from that selected during library preparation (the orange arrows in Figure 1A). When paired-end reads map to the same strand, SVRE assigns them a negative mapping distance. Therefore, a hallmark of inversions is a local mapping distribution that skews towards negative values.

**Figure 1.**
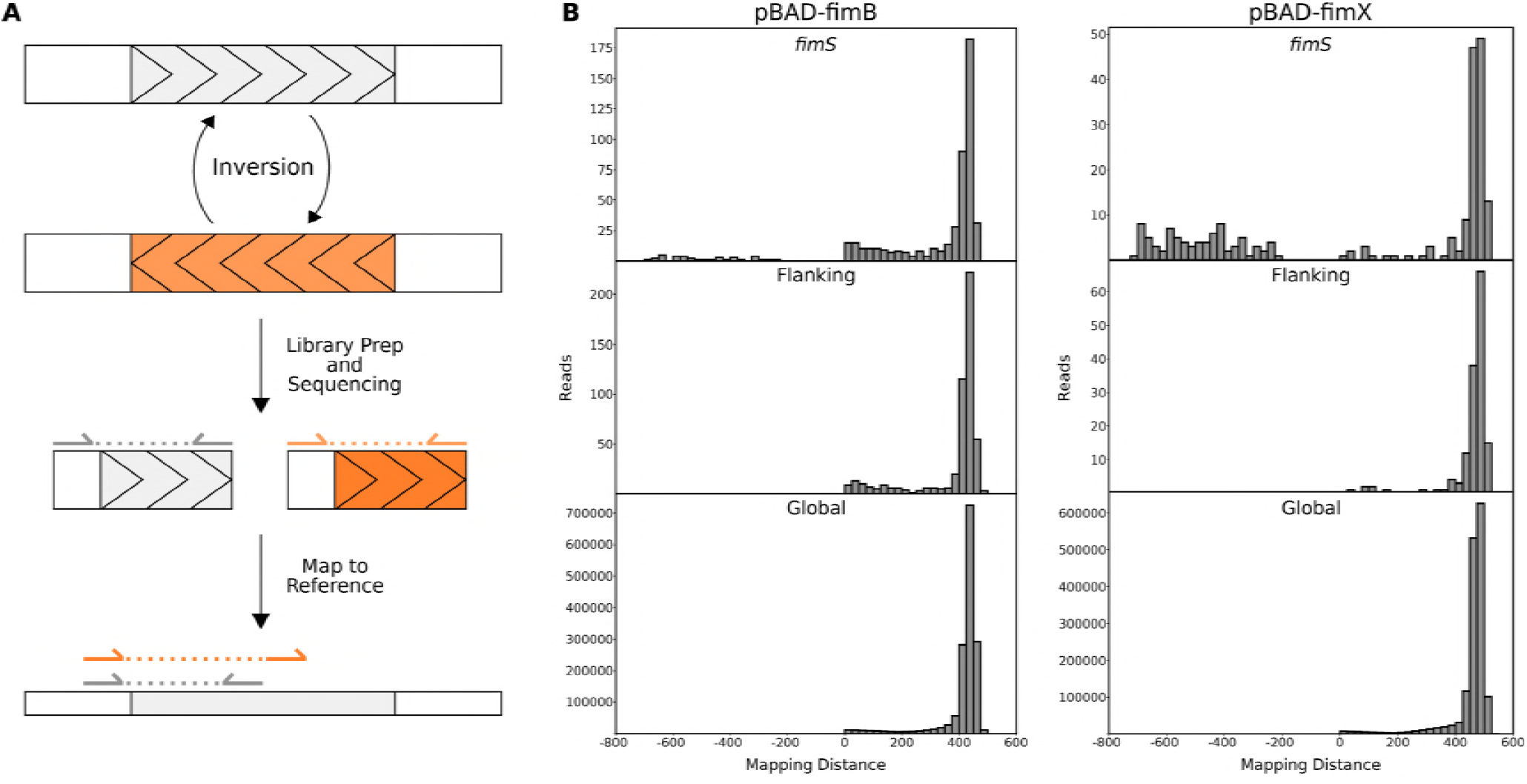
Detection of the *fimS* inversion by the SVRE algorithm. (A) A schematic of how inversions are detected by SVRE. In the right experimental conditions, invertible elements are present in both orientations (shaded gray and orange). After library preparation and sequencing, paired reads derived from sequence in the reference orientation will map to opposite strands of the reference genome with the expected mapping distance. In contrast, paired reads derived from inverted sequences will map to the same strand of the reference genome, resulting in a negative mapping distance, which may also be of an unexpected magnitude. (B) UTI89 carrying a plasmid encoding an arabinose-inducible *fimB* or *fimX* gene was sequenced and analyzed using SVRE. Mapping distance distributions are displayed for windows associated with *fimS* and determined by SVRE to have a significant distribution deviation, windows flanking *fimS*, and the global distribution.

SVRE compares the local mapping distribution of each genome segment to the global distribution, which includes the mapping distances of all paired-end reads genome-wide. The comparison of local and global mapping distributions is made using relative entropy, a statistical test derived from information theory (35). By using relative entropy, SVRE improves on existing SV detection software by providing a more general theoretical foundation for detecting anomalous insert length distributions (as opposed to assuming a normal distribution), resulting in improved signal-to-noise ratio and accuracy. Full theoretical and algorithmic details for SVRE can be found in the Methods and Supplemental Information.

### Application of SVRE to discover SVs in UTI89

SVRE was applied to the uropathogenic strain UTI89 carrying a pBAD33-based plasmid providing arabinose-inducible overexpression of *fimB* or *fimX*, both of which bias the *fimS* switch towards the ON orientation (a similar strategy to that used in (33)). In contrast, the UTI89 reference genome has the *fimS* switch in the OFF orientation; therefore, induction of *fimB* or *fimX* should result in a structural variation (inversion) at *fimS* relative to the published reference sequence. Indeed, with overexpression of either recombinase, windows associated with the *fim* switch showed a local mapping distance distribution that differed from the global distribution (Figure 1B). The difference in the distributions can be primarily attributed to the negative mapping distances observed around the *fim* switch due to paired reads mapping to the same strand, indicative of an inversion. The distribution in flanking windows not associated with *fimS* was similar to the global distribution and these windows were not predicted by SVRE to contain an SV (Figure 1B).

The SVRE algorithm assigns a Relative Information Criterion (RIC) score (i.e. relative entropy) to each window. The RIC score peaks for the *fimS*-associated windows were distinct and well above the genomic background (Figure 2A-B). In addition to the *fimS* peak, there was a distinct peak at *hyxS* in the FimX sample but not the FimB sample. The detection of the *fimS* and *hyxS* peaks with recombinase overexpression demonstrated the ability of SVRE to find known SVs.

**Figure 2.**
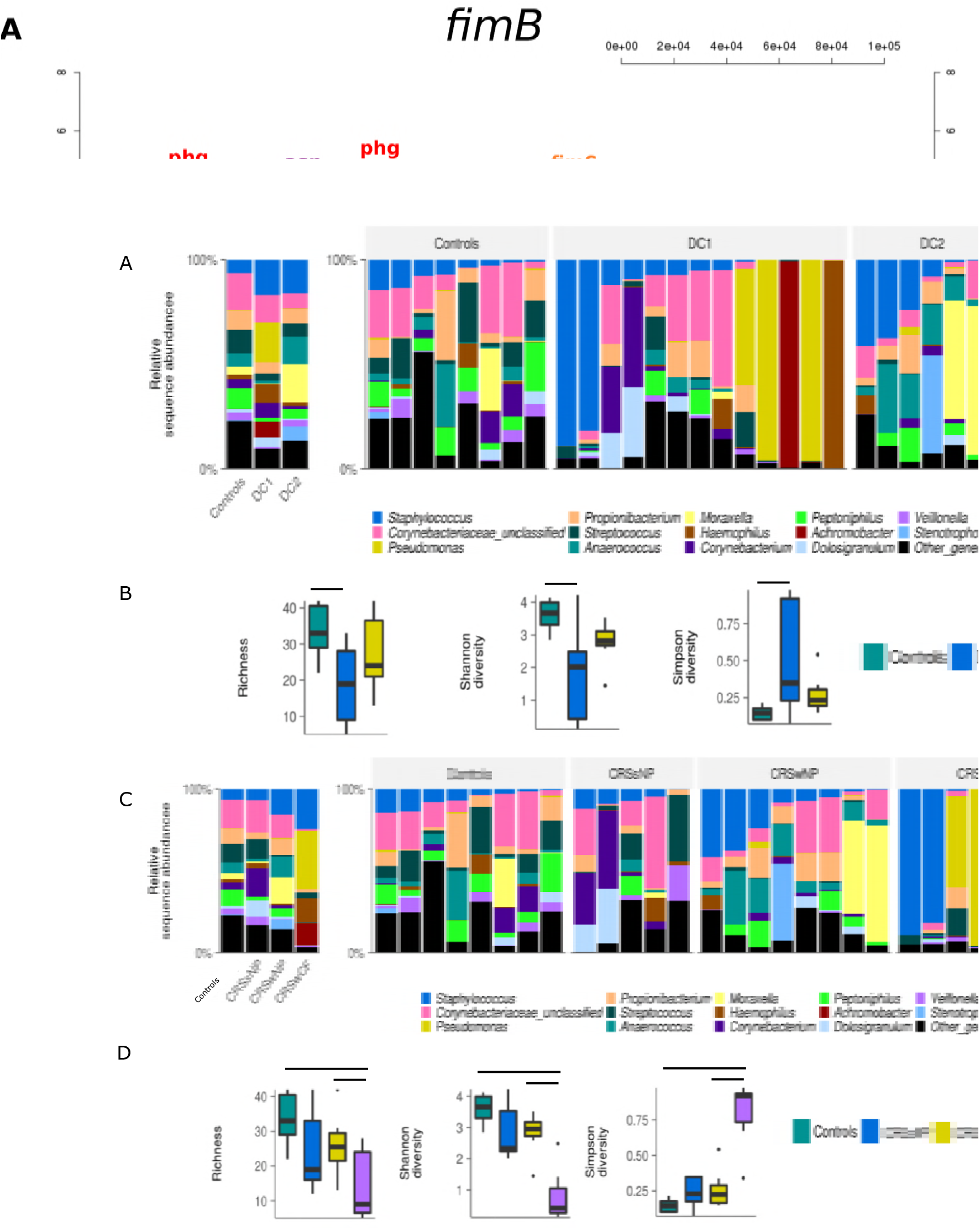
Detection of known and novel structural variations by SVRE in UTI89 overexpressing recombinases. UTI89 cells carrying a plasmid encoding an arabinose-inducible *fimB* (A) or *fimX* (B) gene were sequenced and analyzed using SVRE as in Figure 1. Relative information criterion (RIC) scores are graphed for all windows on the UTI89 chromosome and the pUTI89 plasmid. Peaks are labeled according to the SV they represent as described in the text.

In addition to the *fim* and *hyx* switches, other genomic locations exhibited distinct peaks in RIC scores. Both samples shared a RIC score peak that corresponded to the *ara* locus (labeled “ara” in Figures 2A and B), which is an artefact originating from the use of pBAD plasmids. The remaining peaks included two cases of inversions occurring within prophage (labeled “phg inv” in Figures 2A and B), as well as one inversion occurring in an area containing three asparagine tRNA genes (labeled “asn” in Figures 2A and B). These inversions were predicted to occur in both the FimB and FimX samples. Both samples also shared a prediction of prophage duplication (labeled “dup”), with 2 additional cases of duplication and deletion of prophage (labeled “dup/del”) found only in the FimX sample. Using PCR, each of these SVs was validated in the *fimB* and *fimX* overexpressing strains, but were also found to occur in control cells not overexpressing any recombinases (Figure S1), indicating that these SVs do not appear to be regulated by Fim recombinases. In addition, one of the prophage-associated inversions occurred in the vicinity of a predicted prophage-encoded invertase that is homologous to other phage systems that have been shown to regulate linked prophage promoters (36). The lack of novel invertible elements regulated by FimB and FimX confirms that these recombinases are specific to *fimS* (FimB and FimX) and *hyxS* (FimX).

### Discovery and validation of structural variations in CFT073

The pyelonephritis isolate CFT073 encodes two recombinases (IpuA and IpuB) and one known invertible switch (*ipuS*) that are not found in UTI89 (31). Although IpuB was not able to regulate *fimS*, IpuA was shown to be capable of regulating the *fim* switch both *in vitro* and *in vivo*, adding another layer to Type 1 pili regulation (31). The *ipuS* switch is located between *ipuA* and *ipuR*, and was shown to be inverted by IpuA, IpuB, FimX, and FimE, but not FimB (34).

**Figure 3.**
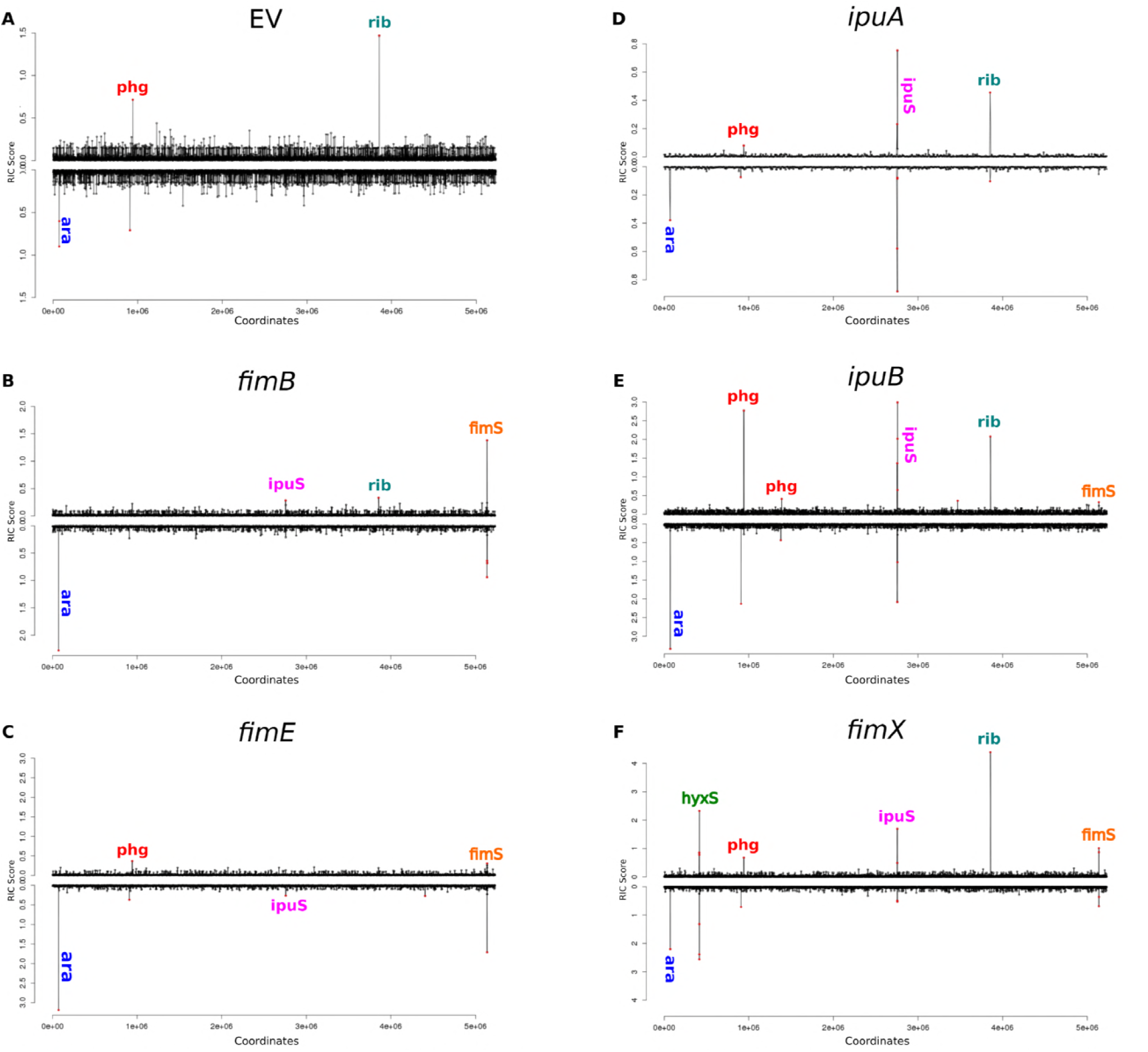
Detection of structural variations using SVRE in CFT073 overexpressing recombinases. Relative information criterion (RIC) scores for all windows on the CFT073 chromosome for (A) cells carrying the pBAD33 control plasmid, or cells overexpressing (B) *fimB*, (C) *fimE*, (D) *ipuA*, (E) *ipuB*, and (F) *fimX*. Significant peaks are labeled according to the SV they represent as described in the text.

The CFT073 allele for each of these recombinases (in cases where they differed from UTI89) was cloned into pBAD33. CFT073 cells carrying each of these plasmids were sequenced and analyzed with SVRE (Figure 3). As expected, a peak for *hyxS* was detected for CFT073/pBAD-fimX cells (Figure 3F), but not for any of the other samples. Distinct peaks for *fimS* were observed for the FimB, FimE, IpuB, and FimX samples (Figure 3B, C, E, F). There were distinct *ipuS* peaks with expression of any of the recombinases (Figure 3B-F). Similar to the UTI89 samples, other peaks were observed that were unrelated to Fim recombinase activity, some of which were present in the empty vector sample (Figure 3A). These included the *ara* operon artefact (“ara” in Figure 3), a false-positive peak associated with mismapping to ambiguous bases in *rrnD* (“rib”), and phage deletions and duplications (“phg”). The phage SVs were found to occur regardless of Fim recombinase expression (Figure S2). Again, as in UTI89, there was no detection of novel invertible elements regulated by the Fim recombinases.

### Effects of recombinase overexpression on ipuS inversion and expression of neighboring genes

We observed an *ipuS* peak in the pBAD-fimB sample (Figure 3B) despite previous data suggesting that FimB is not able to invert *ipuS* (34). To investigate this further, *ipuS* in the ON and OFF orientation was cloned onto a pUC19 backbone. The plasmid sequences confirmed the seven-nucleotide IRs that were observed previously (Figure 4A) (34). Each recombinase was expressed in the MDS42 strain background (chosen due to its lack of endogenous recombinases) in the presence of the *ipuS*-OFF or *ipuS*-ON plasmids (Figure 4B). FimB was capable of inverting *ipuS*, but it had the lowest efficiency of all the recombinases (Figure 4B). The ability of FimB to invert *ipuS* was confirmed in CFT073 (Figure 4C). Overall, IpuB and FimE exhibited the greatest efficiency in OFF to ON inversion, whereas IpuA was most efficient at ON to OFF inversion (Figure 4B-C). These data demonstrate that all of the recombinases, including FimB, are capable of facilitating the inversion of *ipuS*, further validating the accuracy of the SVRE predictions.

**Figure 4.**
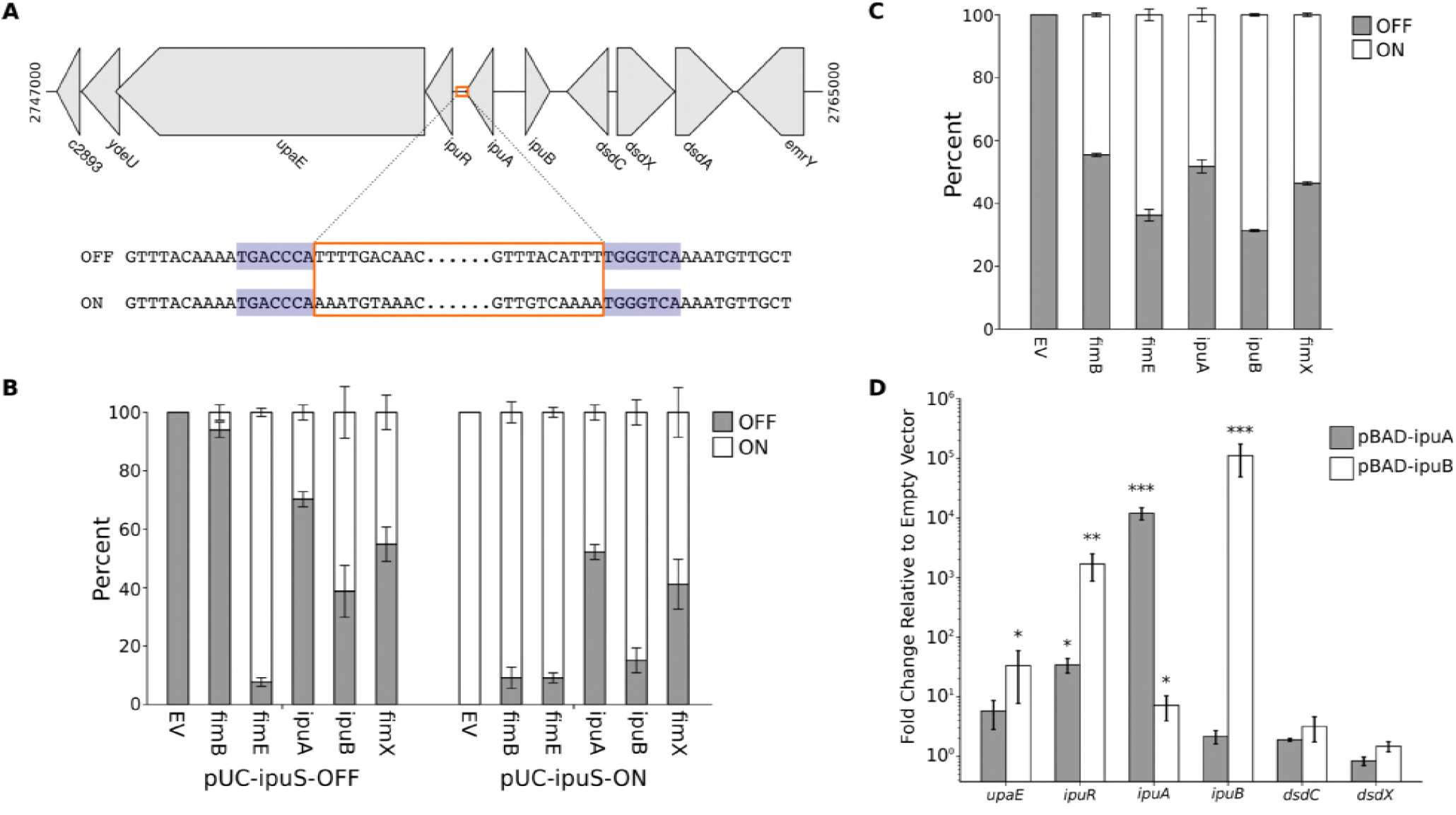
The *ipuS* switch can be inverted by any of the Fim recombinases to drive expression of *ipuR* and *upaE*. (A) A schematic of the genomic location of the *ipuS* invertible element, with *ipuS* outlined in orange, and the 7 bp IRs highlighted in blue. The breakpoints were determined by cloning the invertible element and surrounding sequence from CFT073/pBAD-ipuA induced with arabinose, followed by Sanger sequencing. (B) Quantification of *ipuS* orientation in MDS42 carrying pSLC-372, which contains *ipuS* in the OFF orientation, or pSLC-373, which contains *ipuS* in the ON orientation. The cells also carry a plasmid encoding one of the recombinases or an empty vector control (“EV”). Orientation was quantified via PCR to amplify across the switch, followed by PacI digestion, and measurement of band density using ImageJ. (C) The orientation of the *ipuS* switch was quantified as in B in WT CFT073 with induced expression of different recombinases (“EV” is the empty vector control). (D) CFT073 carrying pBAD33, pBAD-fimE, or pBAD-fimX were induced with arabinose and RT-qPCR was performed to quantify relative gene expression. Gene expression was normalized to 16S levels, and the expression levels are expressed relative to the pBAD33 control samples. The ACt values of each condition were compared to that of the pBAD33 sample using an unpaired, two-tailed T test. * P < 0.05, ** P < 0.01, *** P < 0.001. For figures B-D, bars indicate the mean with error bars representing the standard error of the mean.

It was previously demonstrated that the orientation of the *ipuS* switch can regulate expression of *ipuR* and *upaE* (34). It has also been hypothesized that IpuA may regulate expression of the D-serine utilization locus (37). To delineate the genes that are affected by *ipuS* inversion, RT-qPCR was used to quantify relative expression of several genes in CFT073 cells overexpressing IpuA or IpuB (Figure 4D). No significant change of expression was observed for *dsdC* or *dsdX*, indicating that neither IpuA, IpuB, nor the orientation of *ipuS* affect expression of the D-serine utilization locus. In contrast, expression of *ipuR* was increased by ~1600-fold with IpuB overexpression, and ~34-fold with IpuA overexpression (Figure 4D); this correlates with the orientation of the *ipuS* promoter switch. The significant increase in *upaE* expression was not as dramatic, ~33-fold with IpuB overexpression. Together, these data suggest that *ipuS* inversion only affects the expression of *ipuR* and *upaE* and clarifies that *dsdC* and *dsdX* transcription are not controlled by *ipuS*.

## Discussion

The *fimS* switch is a well-studied example of epigenetic regulation by DNA inversion (29, 38, 39). A single bacterium can give rise to two populations which differ only in the orientation of the *fimS* switch, and individual bacteria can convert between these two populations. The inversion of this switch was first noted to be controlled by two linked recombinases, FimB and FimE (30); in general, *fimS* inversion is described as stochastic, though regulation of the recombinases and several other proteins which bind to regions in the *fimS* switch can influence the bias (15, 38). Therefore, Type 1 pilus expression exhibits phase variation (stochastic inversion) that is responsive to environmental conditions (regulation of bias). With the sequencing of the genomes of several UPEC strains, most notably CFT073 (40) and UTI89 (41), genes encoding additional recombinases with homology to FimB and FimE were discovered (31, 32). These recombinases, like FimB and FimE, were found to regulate inversion of promoter elements genetically linked to the respective recombinase gene. Interestingly, these recombinases also have activity at *fimS*, providing potentially additional layers of regulation for Type 1 pilus expression (31, 32). Importantly, the inverted repeats for these known switches do not always share obvious sequence similarity (see below), implying that a simple search for similar inverted sequences in the genome is not a viable strategy for discovering other invertible switches. The discovery of these unlinked recombinases, therefore, raises several salient questions: (i) do the *fim*-linked FimB and FimE recombinases also have other inversion targets in the genome; (ii) what is the full suite of targets for all of the Fim recombinases; (iii) what is the consequence of coordinating inversion of multiple promoters with the same recombinases; (iv) are the other *non-fim* promoters important for Type 1 pilus expression or function; (v) what additional control of Type 1 pilus expression, if any, is gained by using an unlinked recombinase instead of or in addition to regulating FimB and FimE; (vi) is the regulation of the *fimS* switch important for the evolution or maintenance of the unlinked recombinases, particularly since they are not conserved in all *E. coli* (and thought to be on at least partially mobile elements). We have used whole genome sequencing, combined with overexpression of individual recombinases, to answer the first two of these questions. We found that the *fim* recombinases are very specific, and at least for CFT073 and UTI89, there are no other inversion targets for any of the recombinases aside from those already known. This therefore limits the complexity of questions (iii) and (iv) above, while further shedding light on question (vi) regarding the importance of Type 1 pili and its regulation in *E. coli*.

Positive verification of a new inversion locus is relatively straightforward once the locus is known, and two recent studies have used whole genome sequencing (with Illumina and PacBio data) to achieve accurate quantification of *fimS* inversion percentages under different conditions (42, 43). However, to truly establish the specificity of the *fim* recombinases, a strong negative predictive value is required when analyzing whole genome sequencing data (alternatively, a low noise level). With SVRE, we have improved the analysis of insert read lengths from paired-end short read sequencing data, leading to both sensitive and specific detection of inversions throughout the genome. The key analytical contribution of SVRE is to apply a theoretically optimal measure of differences in distributions (from an information theory perspective) that can then be related to the underlying structure of the genome. More explicitly, currently popular second-generation sequencing technology generates paired-end reads; the reads within each pair are separated by a certain distance, determined by the library preparation. Importantly, the distribution of distances should not depend on the DNA sequence itself (or location on the genome). Therefore, we can use a comparison of local versus global insert length distributions to identify when the genome structure does not match our expectation. This type of analysis is also referred to as anomaly detection, in which relative entropy is a commonly used technique (44). Many other SV detection programs use the same underlying idea, in which anomalous insert lengths are equated to variation in the genome structure, but they make the assumption that the read length distribution is normal (45, 46). Our use of relative entropy in SVRE therefore brings several key advantages: (i) generality to any distribution of insert lengths (which may change depending on how library preparation and size selection are done); (ii) elimination of parameters required to tune the program (such as specifying the expected mean and variance of the assumed normal distribution); (iii) utilization of information contained in “concordant” reads that are within the bulk of the expected distribution (these are still used in the calculation of relative entropy); and (iv) removal of the need for a cutoff for number of “discordant” reads.

From a practical point of view, we find that SVRE produces generally low background signals for most of the genome, from which known SVs clearly stand out (Figure 2A and 2B, between 3.5-4.5 Mbp). To make an assessment of the value of using information theory to analyze read length distributions, we reanalyzed our sequencing data with five other commonly used programs including GASVPro (47), SVDetect (46), Pindel (48), breseq (49), and DELLY (45) (Figure S3). In general, DELLY showed the greatest agreement with SVRE, while GASVPro had the least overlap. Some of these algorithms, such as GASVPro and Pindel, produced many more predictions than SVRE, and required applying a cutoff to allele depth in order reduce the calls to a manageable number. A clear advantage of SVRE is that it enables a simple visualization of the relative entropy (Figures 2 and 3), in addition to providing a list of SV predictions. The connection between DNA structure and relative entropy provides a natural priority ranking for validation and study of individual SVs. Use of SVRE on UTI89 and CFT073 thus allowed us to identify all previously known targets of the Fim recombinases as invertible sequences in the genome. We also identified several SVs that were unrelated to the Fim recombinases. Finally, the good signal-to-noise ratio provides confidence that under the conditions tested, we indeed found no additional invertible elements in the entire genome.

Among the previously identified inversion loci, we found that *ipuS* could be inverted by FimB, both in its native context in the CFT073 chromosome (Figure 3) and when the *ipuS* switch was inserted into a plasmid (Figure 4). In contrast, the original work identifying *ipuS* concluded that FimB was not capable of inverting *ipuS* (34). We did find that, of the five Fim recombinases, FimB inverted *ipuS* in either direction with the lowest efficiency (Figure 4B-C), making its effects more difficult to detect. Combined with differences in the chosen promoters to drive FimB expression, this possibly accounts for the discrepancy between the two studies. Our results also confirm that *ipuS* orientation regulates expression of *ipuR* and *upaE*, while clarifying that the *dsd* operon is not regulated by *ipuS* (Figure 4D). Interestingly, FimE strongly drove inversion from OFF to ON in the MDS42 background (Figure 4B) but not in the CFT073 background (Figure 4C). Of note, while traditionally FimE was thought to only mediate inversion in the ON to OFF direction, FimE has been noted to mediate OFF to ON inversion in some conditions in different strains (42, 50). Therefore, these FimE results could be due to the allele of FimE or other strain-dependent differences.

It is remarkable that Type 1 pilus expression is regulated by five Fim recombinases that regulate little else. The convergence at *fimS* suggests a potentially intricate coordination to control Type 1 pili expression; presumably this facilitates optimal host colonization or adhesion in some other evolutionarily relevant environment. The genetic context for these recombinases may provide some hints as to how *fimS* regulation by both “core” and “accessory” recombinases has evolved. FimB and FimE are considered to be core recombinases since they are encoded adjacent to *fimS* and are present in nearly all *E. coli* strains (51). In contrast, the accessory recombinases FimX, IpuA, and IpuB are encoded at distal locations on two different pathogenicity islands. FimX is encoded adjacent to *hyxS*, while IpuA and IpuB are encoded adjacent to *ipuS*. Therefore, it seems likely that the original role of FimX was to regulate *hyxS*, while IpuA and IpuB originally regulated *ipuS*. We speculate that once UPEC acquired the pathogenicity islands housing these recombinases, the recombinases were co-opted to regulate *fimS* in addition to their cognate switch, and that this additional layer of regulation has given UPEC some sort of advantage. This idea is supported by the observation that *fimX* is enriched in UPEC strains (83.2%) compared to commensals (36%) (51). However, *ipuA* and *ipuB* are found at low levels in roughly equal proportions among UPEC (23.7%) and commensals (15%) alike (51). How these three switches, whose IRs differ in length and sequence, could be regulated by multiple recombinases is still not clear and an area for further investigation. FimB and FimE have been shown to bind to *fimS* at the IRs at half sites that overlap and flank the IRs (52). Therefore, one would hypothesize that the IRs and their surrounding sequence would be quite similar. There is some alignment observed between *ipuS* and *fimS*, and *ipuS* and *hyxS* (34). However, the alignment between *fimS* and *hyxS* is poor, despite the fact that FimX is able to facilitate recombination at both switches (31-33). It thus remains an open question how the Fim recombinases recognize these IRs with apparently dissimilar sequences.

The fact that additional recombinases have been recruited to regulate *fimS* does imply that proper Type 1 pilus expression is important to the evolutionary success of UPEC. This notion is consistent with the observation of positive selection on the FimH adhesin, which results in tuning the conformational flexibility of the protein, leading to modulation of the dynamics of binding to the surface of bladder epithelial cells (53-57). Of note, proper regulation may in some cases include downregulation of Type 1 pili expression at appropriate times, which is also supported by the regulatory mutations seen in EHEC (to lock the *fimS* switch in the OFF orientation) (58), the widespread inactivation of *fimB* in the ST131 *E. coli* lineage via an insertion sequence (42), and the strong positive selection on *fimA* (thought to be due to immune evasion) (59). Downregulation may also explain the finding of low Type 1 pilus expression in bacteria in the urine of some human UTI patients (60-62), though variation in the interaction between different hosts and pathogens during infection is another possibility (63). Here, we have provided additional data that argue that Type 1 pili are important to the success of *E. coli*, and particularly UPEC, suggesting that current efforts to target Type 1 pilus function to prevent and treat UTI represent a rational anti-virulence strategy.

## Materials and Methods

### Bacterial strains

All strains utilized in this study are listed in Table S1. Creation of knockout strains was done using lambda red recombination (64) with 50 bp flanking sequences as described before (65). Primers used for recombination are listed in Table S2.

### Preparation of sequencing data

Overnight cultures were diluted 1:100 into LB broth containing chloramphenicol (20 μg/mL) and were incubated with shaking at 25° C for 24 h, then diluted 1:1000 into fresh media supplemented with chloramphenicol and arabinose (0.5%) and incubated for another 24 h. After the 48 h growth period, genomic DNA was extracted and prepared for Illumina sequencing. For UTI89, the library was prepared using standard techniques including shearing, end-repair, size selection, PCR, and purification with AMPure XP beads; sequencing was performed on an Illumina HiSeq 2000 machine as paired reads with a length of 76 bps. The CFT073 libraries were made using the Illumina TruSeq DNA Library Prep Kit v2 and were sequenced on the Illumina MiSeq as paired reads of a length of 150 bps.

### Development of SVRE

We developed SVRE to improve on existing strategies used in SV detection, particularly those which make use of insert length distributions. When mapped to a perfect reference (i.e. not containing an SV), paired reads will map on opposite strands and at a distance determined by the insert size of the sequencing library, which is usually intentionally controlled during library preparation. Paired reads that map in this way are referred to as “concordant” pairs, while those that do not are “discordant”. One immediate strategy is to focus on discordant reads; clusters of discordant reads mapping to a particular region of the genome are then identified as a potential SV. However, distinguishing between these two classes is not always trivial, and appropriate cutoffs for how many discordant reads should be required to support a true SV are difficult to determine a priori. Programs such as GASVPro (47), SVDetect (46), DELLY (45), VariationHunter (66), BreakDancer (67), and the read distribution module of LUMPY (68) define concordant reads as those whose mapping distances fall within a chosen range based on the expected mapping distance and the standard deviation. In other words, library preparation is assumed to generate a roughly normal distribution of read insert lengths. Another drawback to this approach is that concordant reads are discarded and any information that concordant reads could supply for predicting SVs (such as differences in their length distribution) is lost.

Another strategy that avoids this concordant/discordant differentiation considers the overall distribution of mapping distances. By looking at histograms of mapping distances, changes from the expected distribution can be detected by a number of methods including statistical tests (X^2^, K-S test, t-test, Z-test, etc.) or by using classification algorithms (such as support vector machines). Existing algorithms that utilize this distribution comparison strategy include SVM^2^ (69) and MoDIL (70).

SVRE also uses a distribution comparison strategy. We choose the global insert length distribution as an empirical null model; implicitly, we are assuming that SVs are rare overall and therefore have a minimal global effect on the insert length distribution. We then compare the distribution of a local window to this global distribution using relative entropy (Kullback-Leibler divergence, relative information content, or information divergence/gain). In information theory, relative entropy is a measure of the divergence between two “information” distributions (35). This is strongly related to concepts about signal encoding and compression, in which entropy is known to define an optimal theoretical lower limit for compressed or encoded message size.

With respect to SV detection, to the extent that information is carried within insert length distributions, we suggest that relative entropy is a potentially optimal statistic for quantifying how different a local distribution is from the global null distribution, though we have not formally proven this.

Details about the implementation of SVRE can be found in the Supplemental Information. SVRE was written in Perl and is available for download at https://github.com/swainechen/svre.

### Structural variation prediction with other software

GASVPro version 1.2 (47), SVDetect version 0.8b (46), Pindel version 0.2.5b9 (48), breseq version 0.33.1 (49), and DELLY version 0.7.8 (45) were run according to the instructions provided by the developers. Fastq files were used as the input for breseq, whereas the other programs required sorted, paired-end bam files which were produced using BWA-MEM (71) and SAMtools (72). Any additional pre- and post-processing steps, as well as analysis of the output, were performed ad hoc with Python.

### PCR to confirm structural variations

The primers utilized to validate predicted SVs are listed in Table S2 and were designed according to the specific SV type as outlined in Fig S1A-C. PCR was performed with cells grown for 48 h at 25° C with passaging at 24 h and cells grown for 7 h at 37° C. The cells were grown in LB with arabinose to induce expression of recombinases. PCR was performed with cells from a freshly grown culture or with gDNA isolated from the culture.

### Cloning

The vectors pSLC-372 and pSLC-373 contain the *ipuS* switch in the OFF or ON position, respectively, cloned into the BamHI and SacI sites of pUC19. To obtain *ipuS* DNA in both orientations, *ipuS* was amplified from CFT073/pBAD-ipuA cells induced with arabinose. Plasmids encoding for Fim recombinases were made by amplifying the recombinase from the genomic DNA of either UTI89 or CFT073, and cloning it into the SacI and XbaI sites of pBAD33. The same FimB plasmid was used for both strains given that the *fimB* sequence is identical in the two genomes. These plasmids, along with the primers used for making them, are listed in Table S3.

### Quantification of ipuS orientation

Overnight cultures were diluted 1:100 into 2 mL of LB supplemented with chloramphenicol (20 μg/mL) and arabinose (0.5%) and grown shaking for 7 h at 37° C. A PCR was then performed to amplify across the *ipuS* switch using primers cwr175 and cwr178 to amplify from the genome, or primers M13F and M13R to amplify from the plasmids pSLC-372 and pSLC-373 (Table S2). The resulting product was digested with PacI, which has only one site in the PCR product that is located within *ipuS*. This digestion reaction results in two bands that differ in size depending on the orientation of the switch. The digest reactions were run on a 2% gel, imaged, and the densities of one OFF orientation band and one ON orientation band were quantified using ImageJ FIJI. The total density of the two bands was set to 100% and the percent of ON versus OFF was then calculated.

### RT-qPCR

Overnight cultures of CFT073 carrying pBAD33, pBAD-ipuA, or pBAD-ipuB, were subcultured 1:100 into 10 mL of LB with chloramphenicol (20 μg/mL) in a 100 mL flask and were grown with shaking for 3 h at 37° C. Arabinose was then added to a final concentration of 0.5%, and the cells were allowed to incubate for another hour, at which point 0.5 mL of culture was added to 1 mL of RNAprotect Bacteria Reagent and the cells were lysed using proteinase K and lysozyme. RNA was isolated using the RNeasy Mini Kit, and DNA was removed with DNase I digestion. The SuperScript II RT kit was used to make cDNA. For each sample, a control reaction was run that lacked reverse transcriptase to check for DNA contamination during the qPCR reactions.

Primers employed in the qPCR reaction are listed in Table S2. A control lacking cDNA was included for each pair of primers, in addition to the reactions with and without reverse transcriptase for each sample. The KAPA SYBR FAST qPCR Master Mix was used along with 0.5 μM of each primer and ROX Low. The reactions were run on the ViiA 7 Real-Time PCR System with the following program: 95° C for 3 minutes followed by 40 cycles of 95° C for 3 seconds and 60° C for 20 seconds. The data were analyzed using the ΔΔC_t_ method with 16S acting as a reference gene and the pBAD33 sample as the reference sample. Differences between sample ΔC_t_ values were tested using an unpaired, two-tailed T test.

## Acknowledgments

This work was supported by the National Research Foundation, Singapore (NRF-RF2010-10 to S.L.C.); the National Medical Research Council, Ministry of Health, Singapore grant numbers NMRC/CIRG/1357/2013, NMRC/CIRG/1358/2013, and NMRC/OFIRG/0009/2016; and the Genome Institute of Singapore (GIS) / Agency for Science, Technology, and Research (A*STAR). Experiments were performed by CWR, LTL, BP, SR, and CYC. The SVRE algorithm was developed by RS and SLC. The manuscript was written by CWR, RS, BP, and SLC.

## Supplemental

**Figure S1. Confirmation of novel structural variations in UTI89.** A PCR strategy was employed that was specific to each SV type. (A) For inversions, two sets of primers were used. One set produces a band when the invertible element is in the orientation found on the reference genome. In contrast, the other set produces a band if there is an inversion event. (B) Deletions were detected by using distant primer sets that only produce a band if the intervening sequence is deleted, bringing the priming sites closer together. (C) Duplications were detected using outward facing primer pairs that produce a band only if a tandem duplication event occurs. (D-I) For each SV, the leftmost coordinate of significant windows called by SVRE are represented by red (UTI89/pBAD-fimB) and blue (UTI89/pBAD-fimE) lines. The primers used to confirm the predicted SVs are depicted on the schematic of the neighboring genes, and the gels that resulted from the use of those primers are shown below. (D-F) Confirmation of inversions at (D) 0.9 Mb, (E) 2.1 Mb, and (F) 2.9 Mb were performed in UTI89 (“Ctrl”), UTI89/pBAD33 (“EV”), and UTI89/pBAD-fimX (*“fimX”*) cells. The linked phage invertase *pin* is highlighted in (A). (G-I) Confirmation of (G) a prophage deletion at 1.6 Mb, prophage duplication and deletions at (H) 1.2 Mb and (I) 5.0 Mb. The PCR was performed using WT UTI89 as well as *UTI89ΔfimBΔfimEΔfimX* (“ΔBEX”).

**Figure S2. Confirmation of novel structural variations in CFT073.** For each SV, the leftmost coordinate of significant windows called by SVRE are represented by red (pBAD-fimB), black (pBAD-fimE), orange (pBAD-ipuA), green (pBAD-ipuB), and blue (pBAD-fimX) lines. The primers used to confirm the predicted SVs are depicted on the schematic of the neighboring genes, and the gels that resulted from the use of those primers are shown below. Confirmation of the SVs was performed in CFT073 carrying either pBAD33 (“EV”) or plasmids encoding the various recombinases. (A) Detection of duplication and deletion of phage at 0.9 Mb and (B) a phage at 1.3 Mb.

**Figure S3. Comparison of SVRE calls to that of other SV prediction programs.** SV predictions for (A) UTI89 and (B) CFT073 are listed in the first columns of each table. Whether that SV was detected in a given sample by a program is indicated by a filled box following the color code indicated in the legend.

**Table S1. Strains utilized in this work.** The table lists the strains used in this work. If the strain was part of a previous publication, the appropriate reference is given.

**Table S2. Primers used for strain creation, SV validation, and qRT-PCR.** The table lists primer sets used to detect SVs, create knockout mutant strains, and measure gene expression.

**Table S3. Plasmids utilized in this work.** For each plasmid that was used in this work, either a reference is given or the primers that were used in the creation of the plasmid are listed.

**Supplemental Information. Implementation of SVRE.** A description of how the SVRE program is implemented, including how relative entropy is calculated.

## References

1. Foxman B. 2014. Urinary tract infection syndromes: occurrence, recurrence, bacteriology, risk factors, and disease burden. Infect Dis Clin North Am 28:1–13.

2. Flores-Mireles AL, Walker JN, Caparon M, Hultgren SJ. 2015. Urinary tract infections: epidemiology, mechanisms of infection and treatment options. Nat Rev Microbiol 13:269–84.

3. Foxman B, Barlow R, D’Arcy H, Gillespie B, Sobel JD. 2000. Urinary tract infection: self-reported incidence and associated costs. Ann Epidemiol 10:509–15.

4. Smith AL, Brown J, Wyman JF, Berry A, Newman DK, Stapleton AE. 2018. Treatment and Prevention of Recurrent Lower Urinary Tract Infections in Women: A Rapid Review with Practice Recommendations. J Urol doi:10.1016/j.juro.2018.04.088.

5. Aabenhus R, Hansen MP, Siersma V, Bjerrum L. 2017. Clinical indications for antibiotic use in Danish general practice: results from a nationwide electronic prescription database. Scand J Prim Health Care 35:162–169.

6. Rautakorpi UM, Klaukka T, Honkanen P, Makela M, Nikkarinen T, Palva E, Roine R, Sarkkinen H, Huovinen P, Group MCS. 2001. Antibiotic use by indication: a basis for active antibiotic policy in the community. Scand J Infect Dis 33:920–6.

7. Foxman B. 2010. The epidemiology of urinary tract infection. Nat Rev Urol 7:653–60.

8. Nielubowicz GR, Mobley HL. 2010. Host-pathogen interactions in urinary tract infection. Nat Rev Urol 7:430–41.

9. Bergsten G, Wullt B, Svanborg C. 2005. Escherichia coli, fimbriae, bacterial persistence and host response induction in the human urinary tract. Int J Med Microbiol 295:487–502.

10. Carey AJ, Tan CK, Ipe DS, Sullivan MJ, Cripps AW, Schembri MA, Ulett GC. 2016. Urinary tract infection of mice to model human disease: Practicalities, implications and limitations. Crit Rev Microbiol 42:780–99.

11. Ulett GC, Totsika M, Schaale K, Carey AJ, Sweet MJ, Schembri MA. 2013. Uropathogenic Escherichia coli virulence and innate immune responses during urinary tract infection. Curr Opin Microbiol 16:100–7.

12. Bahrani-Mougeot FK, Buckles EL, Lockatell CV, Hebel JR, Johnson DE, Tang CM, Donnenberg MS. 2002. Type 1 fimbriae and extracellular polysaccharides are preeminent uropathogenic Escherichia coli virulence determinants in the murine urinary tract. Mol Microbiol 45:1079–93.

13. Connell I, Agace W, Klemm P, Schembri M, Marild S, Svanborg C. 1996. Type 1 fimbrial expression enhances Escherichia coli virulence for the urinary tract. Proc Natl Acad Sci US A 93:9827–32.

14. Hultgren SJ, Porter TN, Schaeffer AJ, Duncan JL. 1985. Role of type 1 pili and effects of phase variation on lower urinary tract infections produced by Escherichia coli. Infect Immun 50:370–7.

15. Russell CW, Mulvey MA. 2014. Type 1 and P Pili of Uropathogenic Escherichia coli, p 49–70. In Barocchi MA, Telford JL (ed), Bacterial Pili: Structure, Synthesis and Role in Disease. CAB International, London, UK.

16. Klemm P, Jorgensen BJ, van Die I, de Ree H, Bergmans H. 1985. The fim genes responsible for synthesis of type 1 fimbriae in Escherichia coli, cloning and genetic organization. Mol Gen Genet 199:410–4.

17. Krogfelt KA, Bergmans H, Klemm P. 1990. Direct evidence that the FimH protein is the mannose-specific adhesin of Escherichia coli type 1 fimbriae. Infect Immun 58:1995–8.

18. Zhou G, Mo WJ, Sebbel P, Min G, Neubert TA, Glockshuber R, Wu XR, Sun TT, Kong XP. 2001. Uroplakin Ia is the urothelial receptor for uropathogenic Escherichia coli: evidence from in vitro FimH binding. J Cell Sci 114:4095–103.

19. Eto DS, Jones TA, Sundsbak JL, Mulvey MA. 2007. Integrin-mediated host cell invasion by type 1-piliated uropathogenic Escherichia coli. PLoS Pathog 3:e100.

20. Anderson GG, Palermo JJ, Schilling JD, Roth R, Heuser J, Hultgren SJ. 2003. Intracellular bacterial biofilm-like pods in urinary tract infections. Science 301:105–7.

21. Mulvey MA, Lopez-Boado YS, Wilson CL, Roth R, Parks WC, Heuser J, Hultgren SJ. 1998. Induction and evasion of host defenses by type 1-piliated uropathogenic Escherichia coli. Science 282:1494–7.

22. Martinez JJ, Mulvey MA, Schilling JD, Pinkner JS, Hultgren SJ. 2000. Type 1 pilus-mediated bacterial invasion of bladder epithelial cells. EMBO J 19:2803–12.

23. Wright KJ, Seed PC, Hultgren SJ. 2007. Development of intracellular bacterial communities of uropathogenic Escherichia coli depends on type 1 pili. Cell Microbiol 9:2230–41.

24. Mulvey MA, Schilling JD, Hultgren SJ. 2001. Establishment of a persistent Escherichia coli reservoir during the acute phase of a bladder infection. Infect Immun 69:4572–9.

25. Justice SS, Hung C, Theriot JA, Fletcher DA, Anderson GG, Footer MJ, Hultgren SJ. 2004. Differentiation and developmental pathways of uropathogenic Escherichia coli in urinary tract pathogenesis. Proc Natl Acad Sci U S A 101:1333–8.

26. Spaulding CN, Klein RD, Schreiber HLt, Janetka JW, Hultgren SJ. 2018. Precision antimicrobial therapeutics: the path of least resistance? NPJ Biofilms Microbiomes 4:4.

27. Mydock-McGrane LK, Hannan TJ, Janetka JW. 2017. Rational design strategies for FimH antagonists: new drugs on the horizon for urinary tract infection and Crohn’s disease. Expert Opin Drug Discov 12:711–731.

28. Olsen PB, Klemm P. 1994. Localization of promoters in the fim gene cluster and the effect of H-NS on the transcription of fimB and fimE. FEMS Microbiol Lett 116:95–100.

29. Abraham JM, Freitag CS, Clements JR, Eisenstein BI. 1985. An invertible element of DNA controls phase variation of type 1 fimbriae of Escherichia coli. Proc Natl Acad Sci USA 82:5724–7.

30. Klemm P. 1986. Two regulatory fim genes, fimB and fimE, control the phase variation of type 1 fimbriae in Escherichia coli. EMBO J 5:1389–93.

31. Bryan A, Roesch P, Davis L, Moritz R, Pellett S, Welch RA. 2006. Regulation of type 1 fimbriae by unlinked FimB- and FimE-like recombinases in uropathogenic Escherichia coli strain CFT073. Infect Immun 74:1072–83.

32. Xie Y, Yao Y, Kolisnychenko V, Teng CH, Kim KS. 2006. HbiF regulates type 1 fimbriation independently of FimB and FimE. Infect Immun 74:4039–47.

33. Bateman SL, Seed PC. 2012. Epigenetic regulation of the nitrosative stress response and intracellular macrophage survival by extraintestinal pathogenic Escherichia coli. Mol Microbiol 83:908–25.

34. Battaglioli EJ, Goh KGK, Atruktsang TS, Schwartz K, Schembri MA, Welch RA. 2018. Identification and Characterization of a Phase-Variable Element That Regulates the Autotransporter UpaE in Uropathogenic Escherichia coli. MBio 9.

35. Kullback S, Leibler RA. 1951. On Information and Sufficiency. The Annals of Mathematical Statistics 22:79–86.

36. Stern B, Kamp D. 1989. Evolution of the DNA invertase Gin of phage Mu and related site-specific recombination proteins. Protein Seq Data Anal 2:87–91.

37. Anfora AT, Haugen BJ, Roesch P, Redford P, Welch RA. 2007. Roles of serine accumulation and catabolism in the colonization of the murine urinary tract by Escherichia coli CFT073. Infect Immun 75:5298–304.

38. Schwan WR. 2011. Regulation of fim genes in uropathogenic Escherichia coli. World J Clin Infect Dis 1:17–25.

39. Schembri MA, Ussery DW, Workman C, Hasman H, Klemm P. 2002. DNA microarray analysis of fim mutations in Escherichia coli. Mol Genet Genomics 267:721–9.

40. Welch RA, Burland V, Plunkett G, 3rd, Redford P, Roesch P, Rasko D, Buckles EL, Liou SR, Boutin A, Hackett J, Stroud D, Mayhew GF, Rose DJ, Zhou S, Schwartz DC, Perna NT, Mobley HL, Donnenberg MS, Blattner FR. 2002. Extensive mosaic structure revealed by the complete genome sequence of uropathogenic Escherichia coli. Proc Natl Acad Sci U S A 99:17020–4.

41. Chen SL, Hung CS, Xu J, Reigstad CS, Magrini V, Sabo A, Blasiar D, Bieri T, Meyer RR, Ozersky P, Armstrong JR, Fulton RS, Latreille JP, Spieth J, Hooton TM, Mardis ER, Hultgren SJ, Gordon JI. 2006. Identification of genes subject to positive selection in uropathogenic strains of Escherichia coli: a comparative genomics approach. Proc Natl Acad Sci U S A 103:5977–82.

42. Sarkar S, Roberts LW, Phan MD, Tan L, Lo AW, Peters KM, Paterson DL, Upton M, Ulett GC, Beatson SA, Totsika M, Schembri MA. 2016. Comprehensive analysis of type 1 fimbriae regulation in fimB-null strains from the multidrug resistant Escherichia coli ST131 clone. Mol Microbiol 101:1069–87.

43. Zhang H, Susanto TT, Wan Y, Chen SL. 2016. Comprehensive mutagenesis of the fimS promoter regulatory switch reveals novel regulation of type 1 pili in uropathogenic Escherichia coli. Proc Natl Acad Sci U S A 113:4182–7.

44. Oluwasanya PW. 2017. Anomaly Detection: Review and preliminary Entropy method tests. arXiv: 170808813.

45. Rausch T, Zichner T, Schiatti A, Stutz AM, Benes V, Korbel JO. 2012. DELLY: structural variant discovery by integrated paired-end and split-read analysis. Bioinformatics 28:i333–i339.

46. Zeitouni B, Boeva V, Janoueix-Lerosey I, Loeillet S, Legoix-ne P, Nicolas A, Delattre O, Barillot E. 2010. SVDetect: a tool to identify genomic structural variations from paired-end and mate-pair sequencing data. Bioinformatics 26:1895–6.

47. Sindi SS, Onal S, Peng LC, Wu HT, Raphael BJ. 2012. An integrative probabilistic model for identification of structural variation in sequencing data. Genome Biol 13:R22.

48. Ye K, Guo L, Yang X, Lamijer EW, Raine K, Ning Z. 2018. Split-Read Indel and Structural Variant Calling Using PINDEL. Methods Mol Biol 1833:95–105.

49. Deatherage DE, Barrick JE. 2014. Identification of mutations in laboratory-evolved microbes from next-generation sequencing data using breseq. Methods Mol Biol 1151:165–88.

50. Stentebjerg-Olesen B, Chakraborty T, Klemm P. 2000. FimE-catalyzed off-to-on inversion of the type 1 fimbrial phase switch and insertion sequence recruitment in an Escherichia coli K-12 fimB strain. FEMS Microbiol Lett 182:319–25.

51. Bateman SL, Stapleton AE, Stamm WE, Hooton TM, Seed PC. 2013. The type 1 pili regulator gene fimX and pathogenicity island PAI-X as molecular markers of uropathogenic Escherichia coli. Microbiology 159:1606–17.

52. Gaily DL, Leathart J, Blomfield IC. 1996. Interaction of FimB and FimE with the fim switch that controls the phase variation of type 1 fimbriae in Escherichia coli K-12. Mol Microbiol 21:725–38.

53. Sokurenko EV, Chesnokova V, Dykhuizen DE, Ofek I, Wu XR, Krogfelt KA, Struve C, Schembri MA, Hasty DL. 1998. Pathogenic adaptation of Escherichia coli by natural variation of the FimH adhesin. Proc Natl Acad Sci U S A 95:8922–6.

54. Weissman SJ, Beskhlebnaya V, Chesnokova V, Chattopadhyay S, Stamm WE, Hooton TM, Sokurenko EV. 2007. Differential stability and trade-off effects of pathoadaptive mutations in the Escherichia coli FimH adhesin. Infect Immun 75:3548–55.

55. Chen SL, Hung CS, Pinkner JS, Walker JN, Cusumano CK, Li Z, Bouckaert J, Gordon JI, Hultgren SJ. 2009. Positive selection identifies an in vivo role for FimH during urinary tract infection in addition to mannose binding. Proc Natl Acad Sci U S A 106:22439–44.

56. Schwartz DJ, Kalas V, Pinkner JS, Chen SL, Spaulding CN, Dodson KW, Hultgren SJ. 2013. Positively selected FimH residues enhance virulence during urinary tract infection by altering FimH conformation. Proc Natl Acad Sci U S A 110:15530–7.

57. Kalas V, Pinkner JS, Hannan TJ, Hibbing ME, Dodson KW, Holehouse AS, Zhang H, Tolia NH, Gross ML, Pappu RV, Janetka J, Hultgren SJ. 2017. Evolutionary fine-tuning of conformational ensembles in FimH during host-pathogen interactions. Sci Adv 3:e1601944.

58. Roe AJ, Currie C, Smith DG, Gally DL. 2001. Analysis of type 1 fimbriae expression in verotoxigenic Escherichia coli: a comparison between serotypes O157 and O26. Microbiology 147:145–52.

59. Boyd EF, Hartl DL. 1998. Diversifying selection governs sequence polymorphism in the major adhesin proteins fimA, papA, and sfaA of Escherichia coli. J Mol Evol 47:258–67.

60. Hagan EC, Lloyd AL, Rasko DA, Faerber GJ, Mobley HL. 2010. Escherichia coli global gene expression in urine from women with urinary tract infection. PLoS Pathog 6:e1001187.

61. Subashchandrabose S, Hazen TH, Brumbaugh AR, Himpsl SD, Smith SN, Ernst RD, Rasko DA, Mobley HL. 2014. Host-specific induction of Escherichia coli fitness genes during human urinary tract infection. Proc Natl Acad Sci U S A 111:18327–32.

62. Bielecki P, Muthukumarasamy U, Eckweiler D, Bielecka A, Pohl S, Schanz A, Niemeyer U, Oumeraci T, von Neuhoff N, Ghigo JM, Haussler S. 2014. In vivo mRNA profiling of uropathogenic Escherichia coli from diverse phylogroups reveals common and group-specific gene expression profiles. MBio 5:e01075–14.

63. Schreiber HLt, Conover MS, Chou WC, Hibbing ME, Manson AL, Dodson KW, Hannan TJ, Roberts PL, Stapleton AE, Hooton TM, Livny J, Earl AM, Hultgren SJ. 2017. Bacterial virulence phenotypes of Escherichia coli and host susceptibility determine risk for urinary tract infections. Sei Transi Med 9.

64. Datsenko KA, Wanner BL. 2000. One-step inactivation of chromosomal genes in Escherichia coli K-12 using PCR products. Proc Natl Acad Sci U S A 97:6640–5.

65. Khetrapal V, Mehershahi K, Rafee S, Chen S, Lim CL, Chen SL. 2015. A set of powerful negative selection systems for unmodified Enterobacteriaceae. Nucleic Acids Res 43:e83.

66. Hormozdiari F, Hajirasouliha I, Dao P, Hach F, Yorukoglu D, Alkan C, Eichler EE, Sahinalp SC. 2010. Next-generation VariationHunter: combinatorial algorithms for transposon insertion discovery. Bioinformatics 26:i350–7.

67. Chen K, Wallis JW, McLellan MD, Larson DE, Kalicki JM, Pohl CS, McGrath SD, Wendl MC, Zhang Q, Locke DP, Shi X, Fulton RS, Ley TJ, Wilson RK, Ding L, Mardis ER. 2009. BreakDancer: an algorithm for high-resolution mapping of genomic structural variation. Nat Methods 6:677–81.

68. Layer RM, Chiang C, Quinlan AR, Hall FM. 2014. LUMPY: a probabilistic framework for structural variant discovery. Genome Biol 15:R84.

69. Chiara M, Pesole G, Horner DS. 2012. SVM(2): an improved paired-end-based tool for the detection of small genomic structural variations using high-throughput single-genome resequencing data. Nucleic Acids Res 40:e145.

70. Lee S, Hormozdiari F, Alkan C, Brudno M. 2009. MoDIL: detecting small indels from clone-end sequencing with mixtures of distributions. Nat Methods 6:473–4.

71. Li H. 2013. Aligning sequence reads, clone sequences and assembly contigs with BWA-MEM. arXiv:13033997v2.

72. Li H, Handsaker B, Wysoker A, Fennell T, Ruan J, Homer N, Marth G, Abecasis G, Durbin R, Genome Project Data Processing S. 2009. The Sequence Alignment/Map format and SAMtools. Bioinformatics 25:2078–9.

